## Supplementary file 1 for "ConNIS and labeling instability: new statistical methods for improving the detection of essential genes in TraDIS libraries"

#### 1 Proofs

##### 1.1 Probability of observing a sequence of empty urns

**Theorem 1.** *Let $S$ be a discrete random variable representing the number of consecutive empty urns in a random sequence of size $n$ that contains $k$ empty urns and $n-k$ filled urns. Let the support of $S$ be $\{1, 2, \ldots, k\}$. Then the probability mass function of $S$ is given by:*

$$\mathbb{P}(S = s) = f(s, n, k) = \frac{\binom{n-s-1}{k-s}}{\binom{n-1}{k-1}}, \tag{1}$$

*where $s$ is the number of empty urns in the sample, and $0 < s \leq k \leq n$.*

*Proof.* For each sequence of empty urns of size $s$, there are two special cases: If the first (or last) $s$ urns are empty, $n-s-1$ urns remain, of which $k-s$ must also be empty. Thus, the number of possible combinations is $\binom{n-s-1}{k-s}$. Note that the $-1$ is necessary because at least one filled urn must be present to the right (or left) of the sequence of empty urns; otherwise, the sequence would exceed the specified length $s$.

Next, consider the remaining $n-s-1$ possible positions where a sequence of $s$ empty urns can start. Each of these non-‘edge cases’ allows for $\binom{n-s-2}{k-s}$ combinations. The $-2$ adjustment is required because at least one filled urn must be present to the left and right of the sequence. Thus, the total number of possible combinations is:

$$2\binom{n-s-1}{k-s} + (n-s-1)\binom{n-s-2}{k-s},$$

which can be rewritten as follows

$$2\binom{n-s-1}{k-s} + (n-s-1)\binom{n-s-2}{k-s} = (n-k+1)\binom{n-s-1}{k-s}. \tag{2}$$

We can show with simple transformations that equation 2 holds:

$$
\begin{aligned}
& 2\binom{n-s-1}{k-s} + (n-s-1)\binom{n-s-2}{k-s} = (n-k+1)\binom{n-s-1}{k-s} \\
\Rightarrow \quad & (n-s-1)\binom{n-s-2}{k-s} = (n-k-1)\binom{n-s-1}{k-s} \\
\Rightarrow \quad & \frac{n-s-1}{n-k-1} = \frac{\frac{(n-s-1)!}{(k-s)!(n-k-1)!}}{\frac{(n-s-2)!}{(k-s)!(n-k-2)!}}.
\end{aligned}
$$

The total number of combinations to choose any $k$ consecutive empty urns from a sequence of $n$ urns is $n-k+1$. Since $n-k+1$ is independent of $s$, the probability of observing a sequence of empty urns of length $s$ given $n$ and $k$ can be expressed as

$$
P(S=s) = f(s,n,k) = \frac{(n-k+1)\binom{n-s-1}{k-s}}{(n-k+1)\sum_{i=1}^{k}\binom{n-i-1}{k-i}} = \frac{\binom{n-s-1}{k-s}}{\sum_{i=1}^{k}\binom{n-i-1}{k-i}}
$$

Next, we show

$$
\sum_{i=1}^{k}\binom{n-i-1}{k-i} = \binom{n-1}{k-1}.
$$

Let $m = n-1$ and by using Pascal's rule and the Hockey-stick identity of Pascal's triangle we get

$$
\begin{aligned}
& \sum_{i=1}^{k}\binom{m-i}{k-i} = \binom{m}{k-1} \\
\Rightarrow \quad & \sum_{i=0}^{k}\binom{m-i}{k-i} = \binom{m+1}{k}
\end{aligned}
$$

Hence, we get the pmf

$$
f(s,n,k) = \frac{\binom{n-s-1}{k-s}}{\binom{n-1}{k-1}}
$$

□

#### 1.2 Convergence in distribution

**Theorem 2.** *Let $(n_m)_{m\in\mathbb{N}}$ and $(k_m)_{m\in\mathbb{N}}$ be sequences of integers such that*

$$
n_m \to \infty, \qquad 1 \le k_m < n_m, \qquad \frac{k_m}{n_m} \to q \in (0,1) \quad \textit{as} \quad m \to \infty.
$$

*Let $p := 1 - q$ and let*

$$g(s,p) = (1-p)^{s-1}p$$

*be the the probability mass function (pmf) of a geometric distribution with $s \in \{1, 2, ...\}$. Further, let $f(s, n_m, k_m)$ be the pmf of Theorem 1. Then, for a random variable $X_m$ with pmfs $f(s, n_m, k_m)$ and a random variable $X$ with pmf $g(s,p)$ we have*

$$X_m \xrightarrow{d} X \quad as \quad m \to \infty.$$

*Proof.* By the Portmanteau Theorem, it suffices to show that $f(s, n_m, k_m)$ converges pointwise to $g(s,p)$:

Fix $s \in \mathbb{N}$. By definition of 1 we have $1 \leq s \leq k_m$. For $m \to \infty$ we have further

$$\begin{aligned} \lim_{m\to\infty} f(s, n_m, k_m) &= \lim_{m\to\infty} \frac{\binom{n_m - s - 1}{k_m - s}}{\binom{n_m - 1}{k_m - 1}} \\ &= \lim_{m\to\infty} \left( \prod_{i=1}^{s-1} \frac{k_m - i}{n_m - i} \cdot \frac{n_m - k_m}{n_m - s} \right). \end{aligned}$$

Setting $q_m := \frac{k_m}{n_m}$ implies $q_m \to q$ and since $s$ is fixed we have

$$\prod_{i=1}^{s-1} \frac{k_m - i}{n_m - i} \xrightarrow[m\to\infty]{} q^{s-1}.$$

Similar, we have

$$\frac{n_m - k_m}{n_m - s} \xrightarrow[m\to\infty]{} 1 - q.$$

Thus,

$$\lim_{m\to\infty} f(s, n_m, k_m) = q^{s-1}(1-q) = (1-p)^{s-1}p = g(s,p).$$

□

#### 1.3 Expectation and Variance

**Theorem 3.** *Given the probability mass function $f(s,n,k)$ of a random variable $S$ as defined in eq.* (1)*, the expected value is given by*

$$\mathbb{E}(S) = \frac{n}{n-k+1},$$

*and the variance by*

$$\mathbb{V}ar(S) = \frac{\binom{n}{k-1} + 2\binom{n}{k-2}}{\binom{n-1}{k-1}} - \frac{\binom{n}{k-1}^2}{\binom{n-1}{k-1}^2}.$$

*Proof.* First, the expectation of the discrete random variable $S$ is

$$\mathbb{E}(S) = \sum_{s=1}^{k} s \frac{\binom{n-s-1}{k-s}}{\binom{n-1}{k-1}}.$$

Since only the numerator depends on $s$, it is sufficient to show

$$\sum_{s=1}^{k} s \binom{n-s-1}{k-s} = \binom{n}{k-1}.$$

We write $s = \binom{s}{1}$ and extend the sum to start at $s = 0$, noting that the term for $s = 0$ is zero. Then

$$\begin{aligned} \sum_{s=1}^{k} s \binom{n-s-1}{k-s} &= \sum_{s=1}^{k} \binom{s}{1} \binom{n-s-1}{k-s} \\ &= \sum_{s=0}^{k} \binom{s}{1} \binom{n-s-1}{k-s}. \end{aligned}$$

Using the symmetry of binomial coefficients,

$$\binom{n-s-1}{k-s} = \binom{n-s-1}{n-k-1},$$

and setting $N := n - 1$, this becomes

$$\sum_{s=0}^{k} \binom{s}{1} \binom{N-s}{N-k}.$$

By a standard Vandermonde-type identity,

$$\sum_{s=0}^{N} \binom{s}{1} \binom{N-s}{N-k} = \binom{N+1}{N-k+1+1} = \binom{n}{k-1},$$

and the extra terms with $s > k$ vanish, so the sum up to $k$ already equals $\binom{n}{k-1}$.

Hence,

$$\mathbb{E}(S) = \frac{1}{\binom{n-1}{k-1}} \sum_{s=1}^{k} s \binom{n-s-1}{k-s} = \frac{\binom{n}{k-1}}{\binom{n-1}{k-1}}.$$

□

Next, we prove the variance using the expectation, thus, for

$$\begin{aligned} \mathbb{V}ar(S) &= \mathbb{E}(S^2) - \mathbb{E}(S)^2 \\ &= \sum_{s=1}^{k} s^2 \frac{\binom{n-s-1}{k-s}}{\binom{n-1}{k-1}} - \left( \sum_{s=1}^{k} s \frac{\binom{n-s-1}{k-s}}{\binom{n-1}{k-1}} \right)^2 \end{aligned}$$

we only need to prove the identity

$$\sum_{s=1}^{k} s^2 \frac{\binom{n-s-1}{k-s}}{\binom{n-1}{k-1}} = \frac{\binom{n}{k-1} + 2\binom{n}{k-2}}{\binom{n-1}{k-1}}.$$

Again, only the numerator is dependent on $s$ so we need to show that

$$\sum_{s=1}^{k} s^2 \binom{n-s-1}{k-s} = \binom{n}{k-1} + 2\binom{n}{k-2}. \tag{3}$$

By expressing $s^2$ as a sum of binomial coefficients and using $\binom{0}{x} = 0$ for $x \in \mathbb{N}_+$ we get

$$\begin{aligned}
\sum_{s=1}^{k} s^2 \binom{n-s-1}{k-s} &= \sum_{s=1}^{k} \left[\binom{s}{1} + 2\binom{s}{2}\right] \binom{n-s-1}{k-s} \\
&= \sum_{s=0}^{k} \left[\binom{s}{1} + 2\binom{s}{2}\right] \binom{n-s-1}{k-s} \\
&= \sum_{s=0}^{k} \binom{s}{1}\binom{n-s-1}{k-s} + 2\sum_{s=0}^{k} \binom{s}{2}\binom{n-s-1}{k-s} \\
&= \binom{n}{k-1} + 2\binom{n}{k-2}
\end{aligned}$$

completing the proof. □

### 2 Extensions of state-of-the-art methods

#### 2.1 Binomial

The *Binomial* approach in the `TSAS 2.0` package [Burger et al., 2017] determines essentiality of a gene $j$ based on the probability to observe a number of IS using a binomial distribution. The approach estimates the success probability of the binomial distribution with the insertion density $\theta$. Our proposed approach weights $\theta$ by the factor $0 < w \leq 1$. Thus, the probability of observing $h_j$ insertion sites under the assumption of non-essentiality is then

$$\mathbb{P}(X = h_j) = f(h_j, b_j, \theta, w) = \binom{b_j}{h_j} (\theta \cdot w)^{h_j} (1 - \theta \cdot w)^{b_j - h_j}. \tag{4}$$

The probability of observing $h_j$ IS by chance will be higher for $w < 1$ compared to $w = 1$ (original method), labeling fewer genes 'essential' for a given significance level.

#### 2.2 Geometric

For this approach, the probability of observing an IS after at least $l_j$ 'failures' (non-insertion sites) is calculated using the geometric distribution. Here, $l_j$ is the longest observed insertion-free sequence in gene $j$, while the genome-wide insertion density $\theta$ is used as success probability for the Bernoulli trials. The modified weighted method is

$$\mathbb{P}(X = l_j) = f(l_j, \theta, w) = (1 - \theta \cdot w)^{l_j}(\theta \cdot w). \tag{5}$$

Lowering $w$ will increase the probability of observing at least $l_j$ insertion-free sites by chance. Thus, for $w < 1$ fewer genes are labeled as 'essential' for a given significance level compared to the original method ($w = 1$).

#### 2.3 Tn5Gaps

The *Tn5Gaps* method of the `Transit` software suite uses the Gumbel distribution to calculate the probability of an insertion-free sequence along the genome [DeJesus et al., 2015]. Therefore, it includes non-coding regions in its analysis and determines the essentiality of a gene $j$ by the insertion-free sequence with the largest overlap $k_j$ in terms of base pairs for gene $j$. Following the definition of the authors, the cumulative distribution function is

$$F(k_j, \mu, \beta) = \exp^{-\exp^{(\mu - k_j)/\beta}}$$

with location parameter $\mu$ and scale parameter $\beta$. These parameters are estimated in the original method based on the genome-wide non-insertion density, i.e., $1-\theta$, and the number of IS over the genome $b$. We weight $\theta$ in the estimators of the location parameters by $w$:

$$\hat{\mu} = \log_{\frac{1}{1-\theta w}}(b \cdot \theta \cdot w)$$

and

$$\hat{\beta} = \frac{1}{\ln(\frac{1}{(1-\theta w)})}.$$

By setting $w < 1$, the estimated values for $\mu$ and $\beta$ will be lower compared to the original method. This increases the probability $1 - F(k_j, \hat{\mu}, \hat{\beta})$ of observing an insertion-free sequence of at least length $k_j$ throughout the genome. Again, compared to the original version fewer genes are labeled as 'essential'.
