## Supplementary file 2 for "ConNIS and labeling instability: new statistical methods for improving the detection of essential genes in TraDIS libraries"

### 1 Results for low and medium dense libraries in semi-synthetic data settings

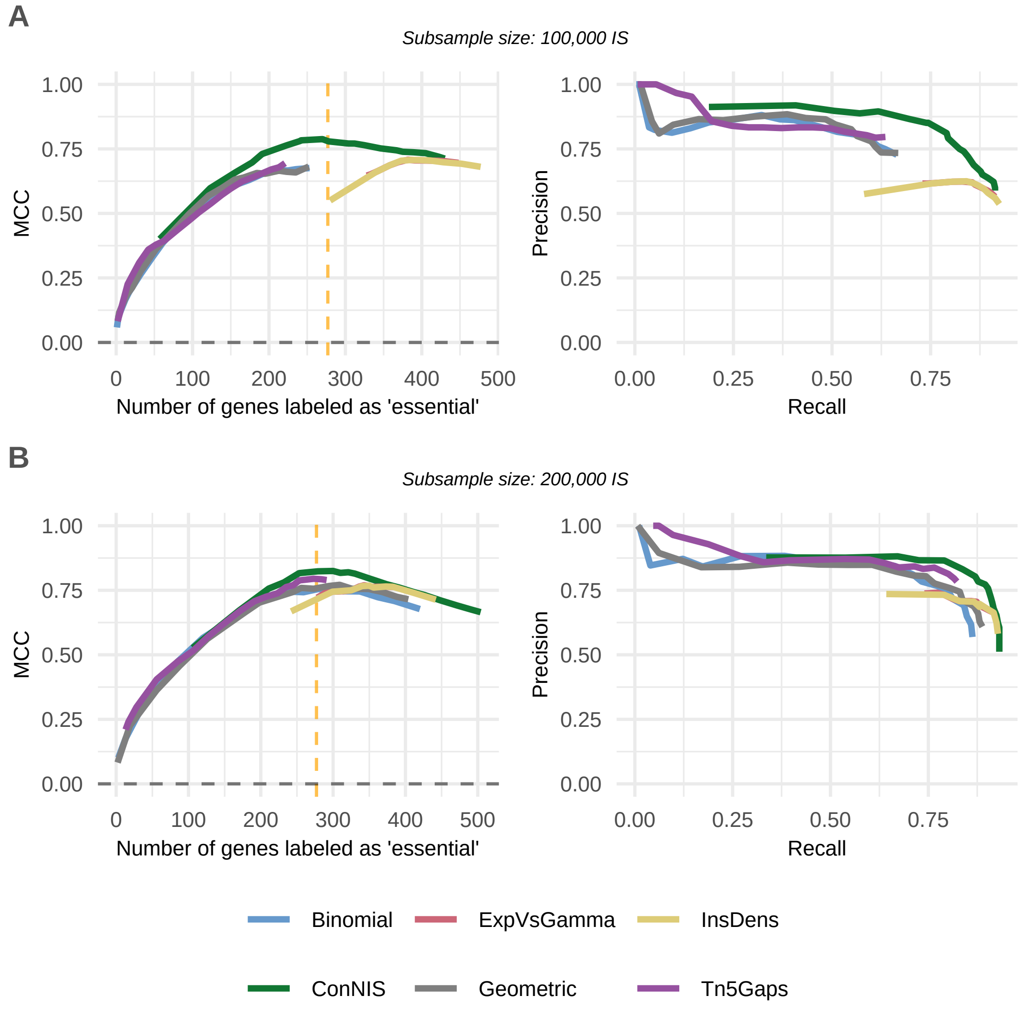


Figure S1: **MCC and PRC performances for semi-synthetic data.** Subsamples were generated by randomly drawing IS from a low- (**A**) and medium-density (**B**) library of *E. coli* BW25113 Goodall et al. (2018). The Kaio library Baba et al. (2006) was used as reference for 'true' gene essentiality. The vertical dotted line shows the true value.

#### 2 Gene-wise insertion densities from real world and simulated data

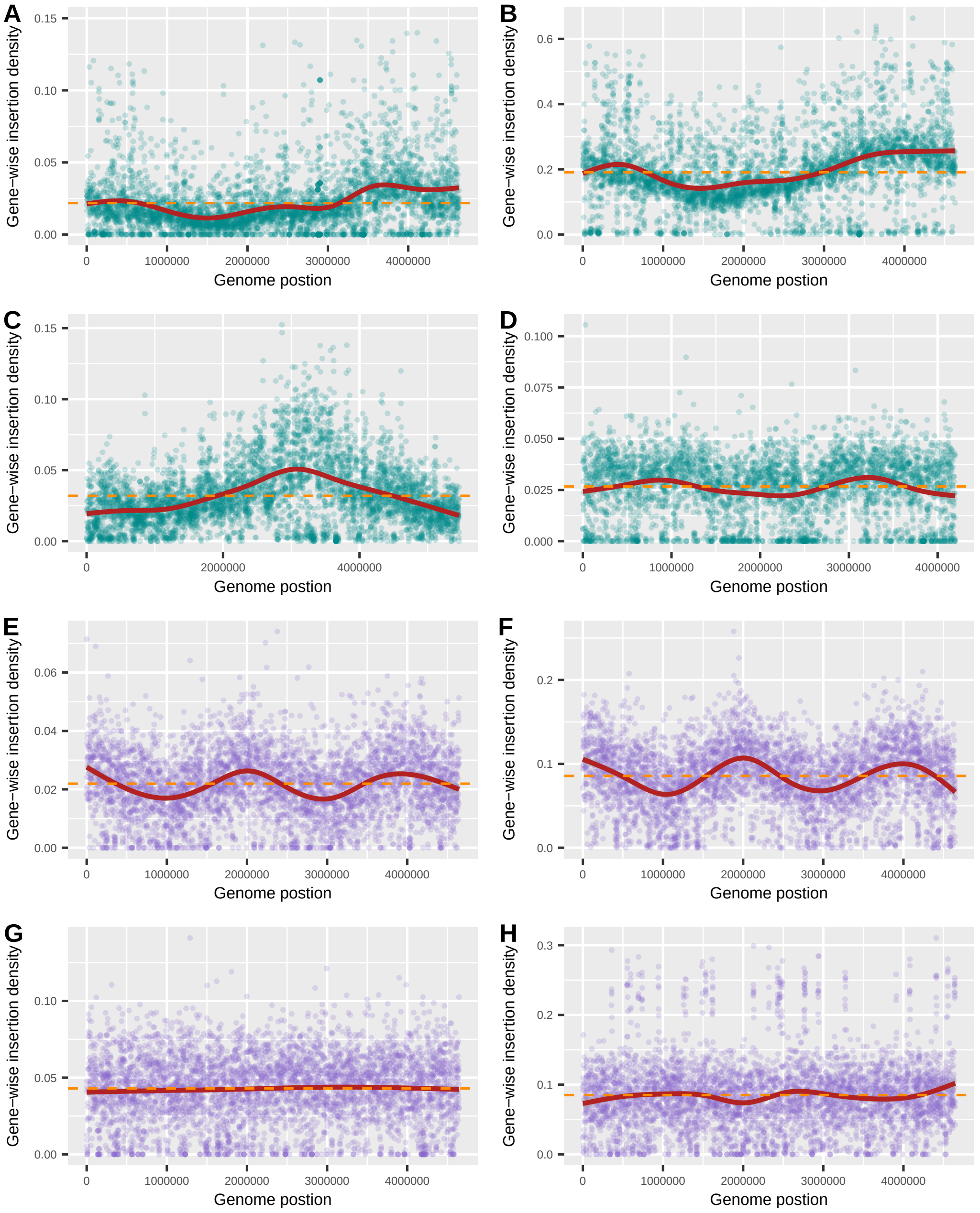


Figure S2: Variability of the distribution of gene-wise insertion densities from real world data (cyan) and our simulated data (purple). The red lines show smoothings of the gene-wise insertion densities a long the genome based on local polynomial regressions while the dashed orange lines show the genome-wide insertion densities. **a)** *E. coli* BW25113 random bar-coded data with ≈100,000 unique insertion sites (Wetmore et al., 2015). **b)** *E. coli* BW25113 mini-Tn5 data with ≈900,000 unique insertion sites Goodall et al. (2018). **c)** *Rhodopseudomonas palustris* CGA009 Tn5-based data with ≈175,000 unique insertion sites (Pechter et al., 2016). **d)** *Sphingobium sp.* SYK-6 Tn5 data with ≈155,000 unique insertion sites (Bleem et al., 2023). **e)** Major synthetic setting 1 with $\pi = 0.9$, wavelength 2,000,000bp, 100,000 unique insertion sites and additional 5000 unique insertions sites as technical noise. **f)** Major synthetic setting 1 with $\pi = 0.95$, wavelength 2,000,000bp, 400,000 unique insertion sites and no technical noise. **g)** Major synthetic setting 2 with uniform essential ORFs with $\pi = 1$, 0 hotspots, 20 coldspots, 200,000 unique insertion sites and additional 5000 unique insertions sites as technical noise. **h)** Major synthetic setting 2 with $\pi = 0.85$, 20 hotspots, 20 coldspots, 100,000 unique insertion sites and additional 5000 unique insertions sites as technical noise.

#### 3 Effect of weight parameter on FNR, FDR and MCC

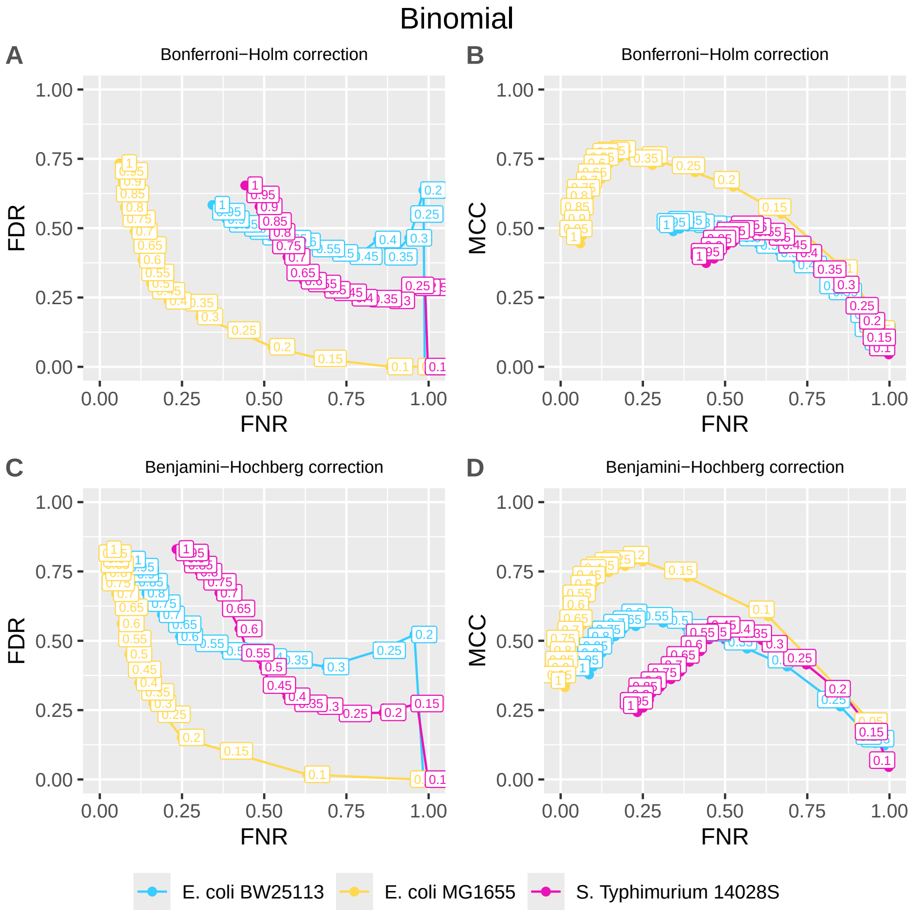


Figure S3: **Effect of weighting the insertion density on the false negative rate (FNR), the false discovery rate (FDR), and MCC for *Binomial*.** The colored numbers are the weighting values $w$ that have been applied with *Binomial* ranging from 0.01 to 1. All results are based on the publicly available IS of two *E. coli* (Wetmore et al., 2015; Ma et al., 2024) and one *S. Typhimurium* (Mandal and Kwon, 2017) real worlds datasets (see Section **??** for details).

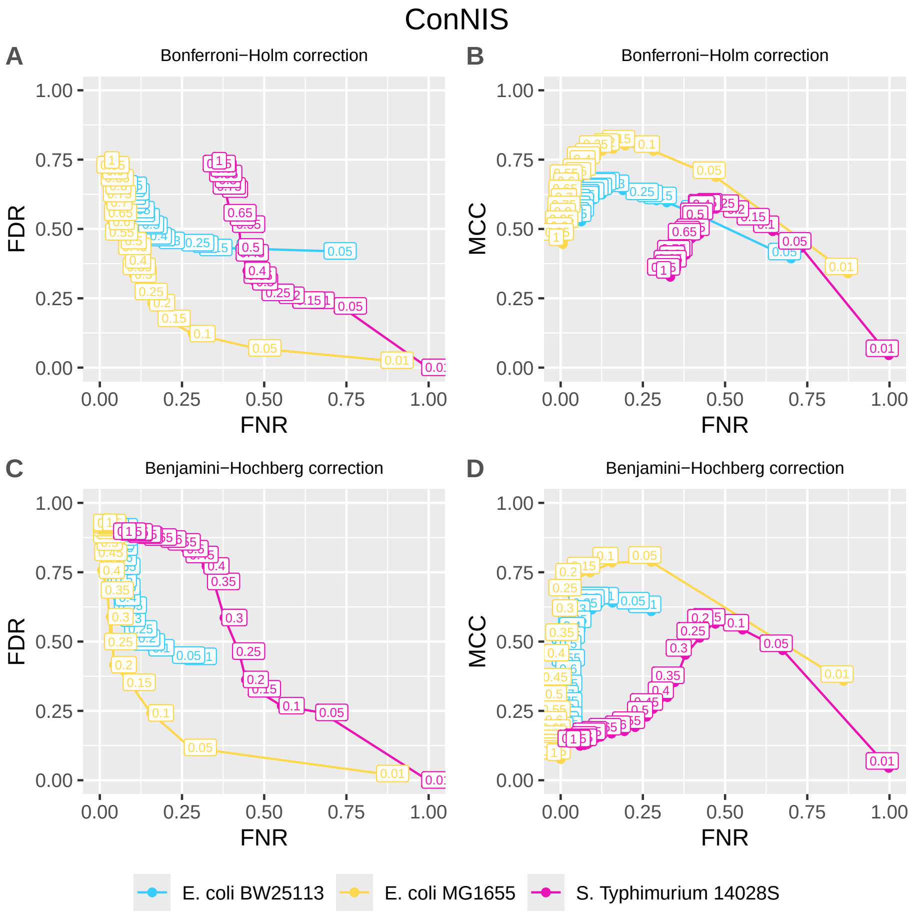


Figure S4: **Effect of weighting the insertion density on the false negative rate (FNR), the false discovery rate (FDR), and MCC for *ConNIS*.** The colored numbers are the weighting values $w$ that were applied with *ConNIS* ranging from 0.01 to 1. All results are based on the publicly available IS of two *E. coli* (Wetmore et al., 2015; Ma et al., 2024) and one *S. Typhimurium* (Mandal and Kwon, 2017) real worlds datasets (see Section **??** for details).

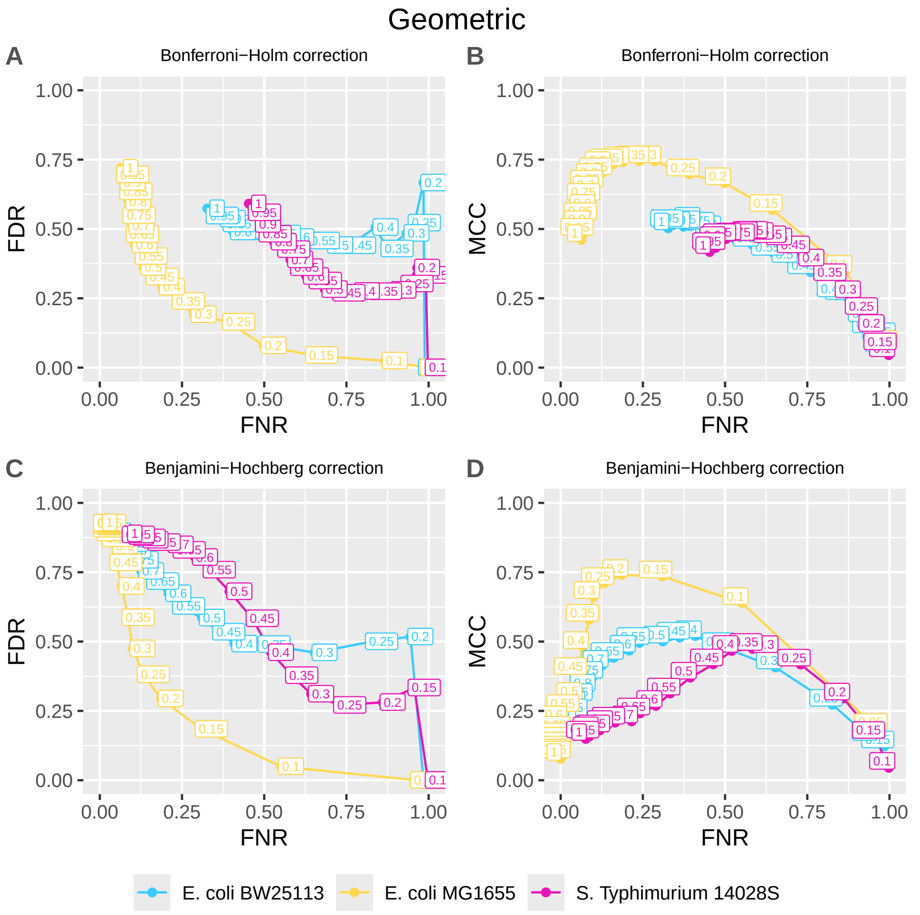


Figure S5: **Effect of the weighting of the insertion density on the false negative rate (FNR), the false discovery rate (FDR), and MCC for *Geometric*.** The colored numbers are the weighting values $w$ that have been applied with *Geometric* ranging from 0.01 to 1. All results are based on the publicly available IS of two *E. coli* (Wetmore et al., 2015; Ma et al., 2024) and one *S. Typhimurium* (Mandal and Kwon, 2017) real worlds datasets (see Section **??** for details).

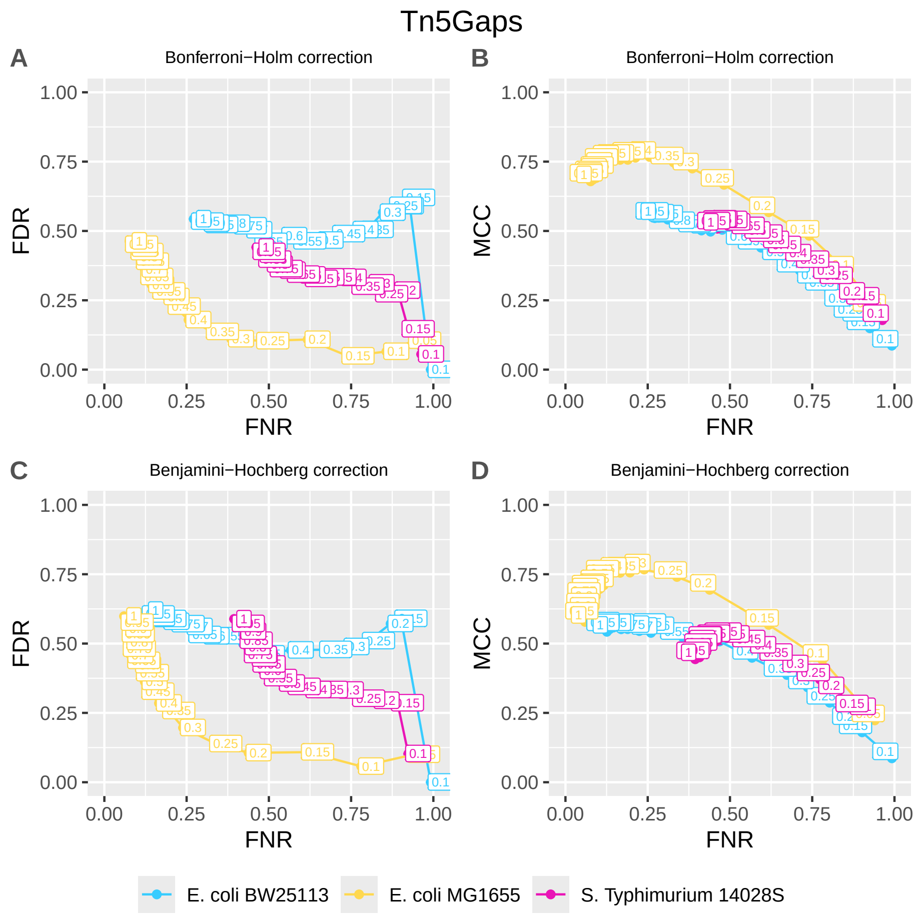


Figure S6: **Effect of weighting the insertion density on the false negative rate (FNR), the false discovery rate (FDR), and MCC for *Tn5Gaps*.** The colored numbers are the weights $w$ that were applied with *Tn5Gaps* ranging from 0.01 to 1. All results are based on the publicly available IS of two *E. coli* (Wetmore et al., 2015; Ma et al., 2024) and one *S. Typhimurium* (Mandal and Kwon, 2017) real worlds datasets (see Section **??** for details).

### 4 Optimal number of selected genes in semi-synthetic setting

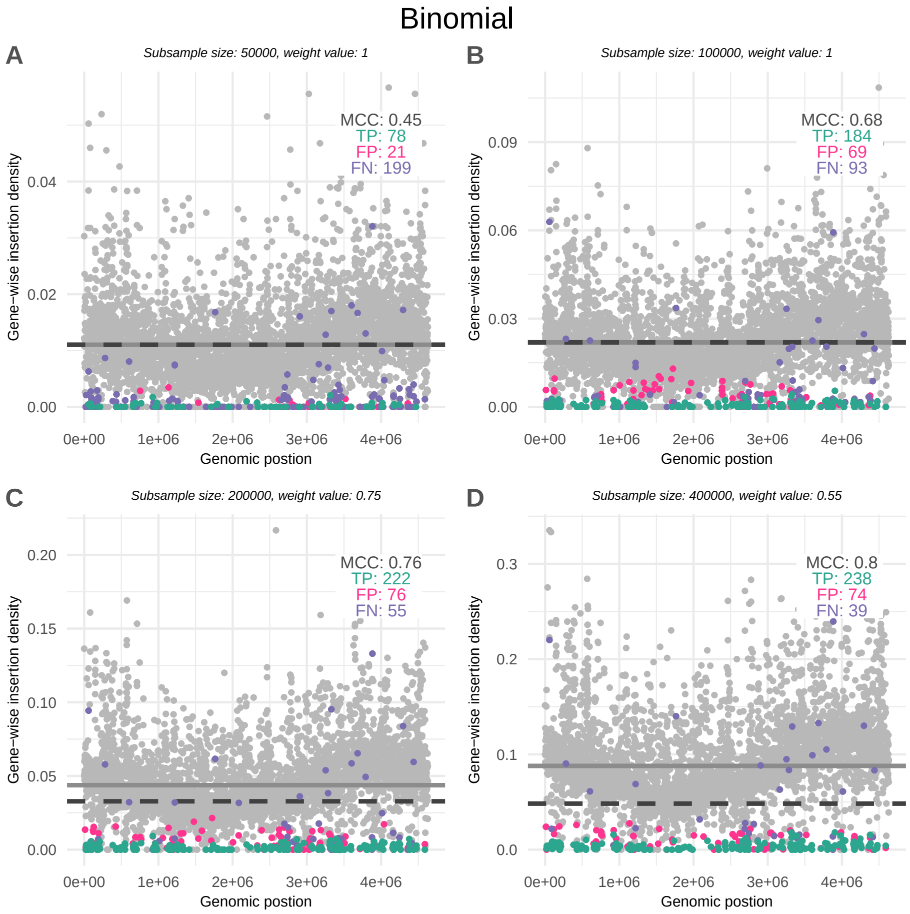


Figure S7: **Selected genes and gene-wise insertion densities for the Binomial approach**. The solid horizontal line shows genome-wide insertion density and the dashed horizontal line the weighted genome-wide insertion density giving the highest MCC. Green, pink and purple dots show the true positives (TP), false positives (FP) and false negatives (FN) selected genes.

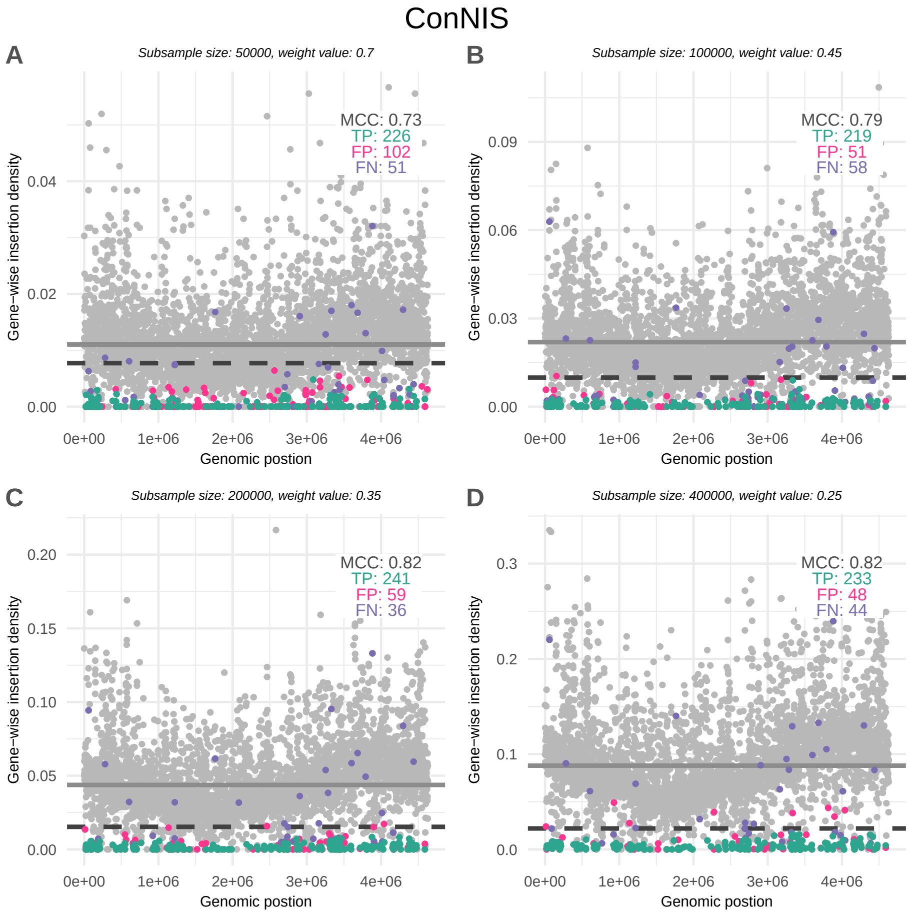


Figure S8: **Selected genes and gene-wise insertion densities for the ConNIS approach**. The solid horizontal line shows genome-wide insertion density and the dashed horizontal line the weighted genome-wide insertion density giving the highest MCC. Green, pink and purple dots show the true positives (TP), false positives (FP) and false negatives (FN) selected genes.

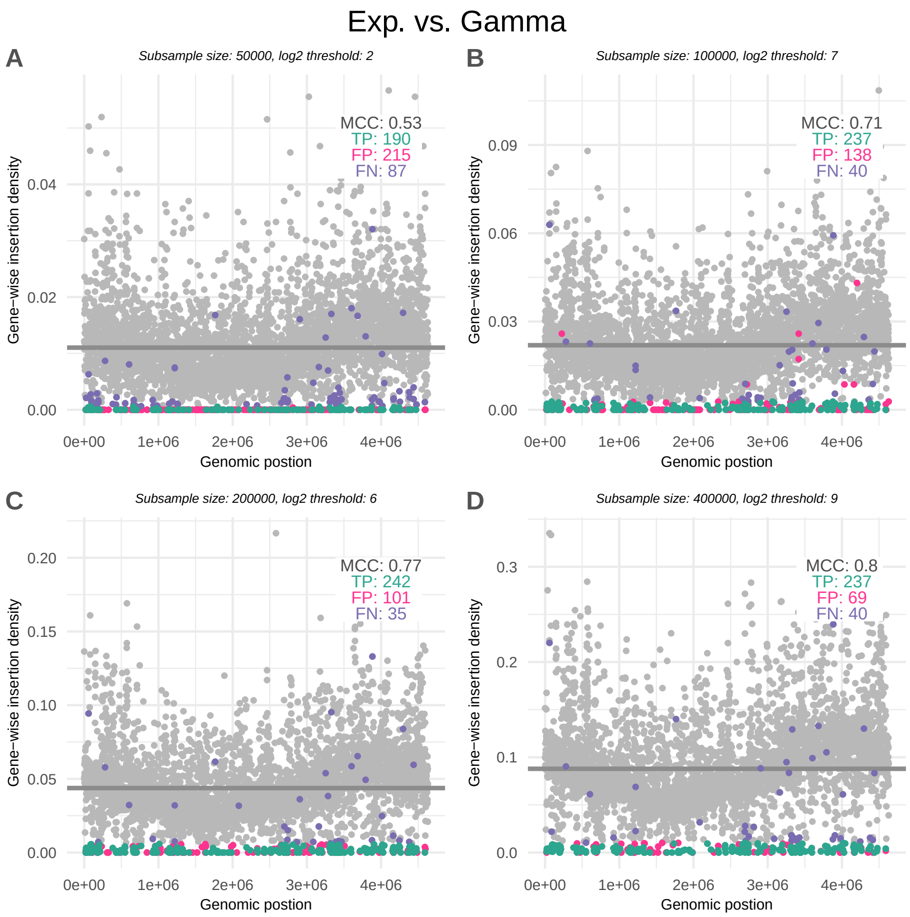


Figure S9: **Selected genes and gene-wise insertion densities for the Exp. vs. Gamma approach**. The solid horizontal line shows genome-wide insertion density. Green, pink and purple dots show the true positives (TP), false positives (FP) and false negatives (FN) selected genes.

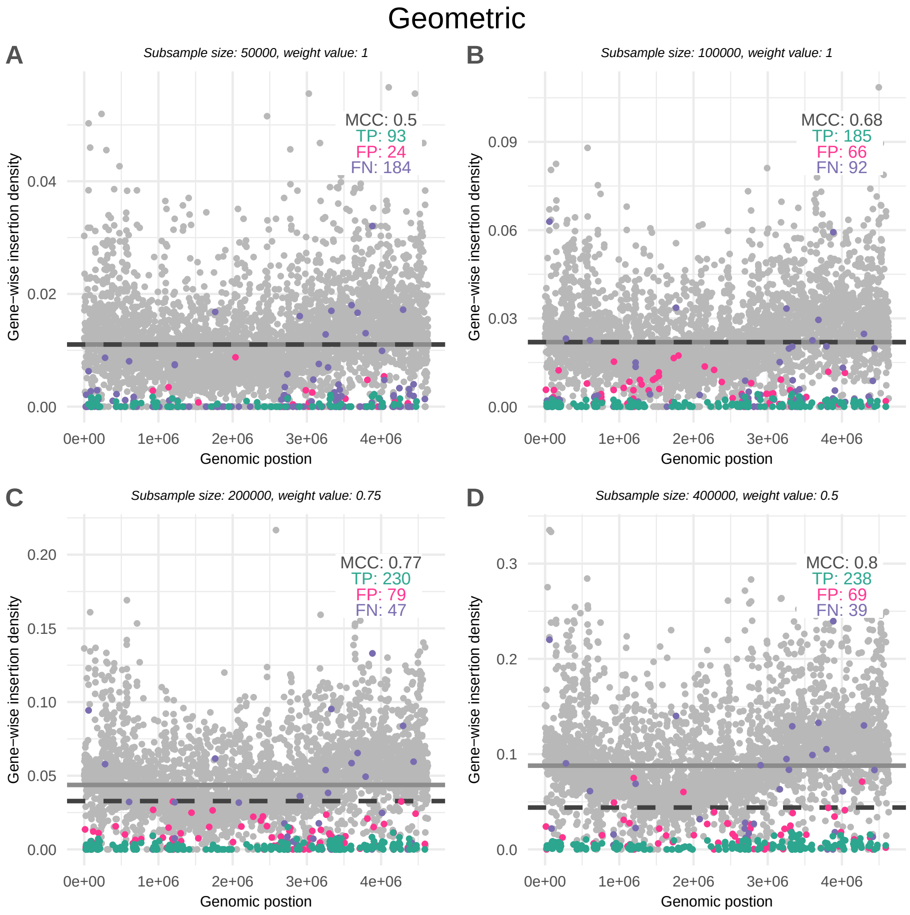


Figure S10: **Selected genes and gene-wise insertion densities for the Geometric approach**. The solid horizontal line shows genome-wide insertion density and the dashed horizontal line the weighted genome-wide insertion density giving the highest MCC. Green, pink and purple dots show the true positives (TP), false positives (FP) and false negatives (FN) selected genes.

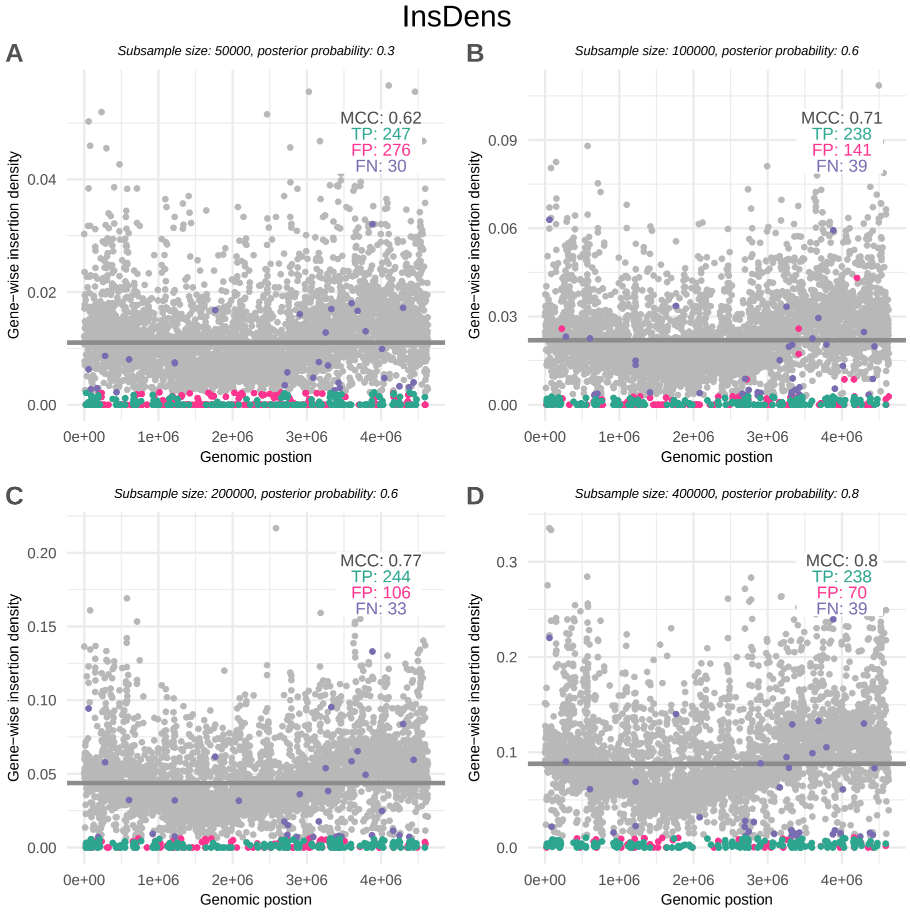


Figure S11: **Selected genes and gene-wise insertion densities for the InsDens approach**. The solid horizontal line shows genome-wide insertion density. Green, pink and purple dots show the true positives (TP), false positives (FP) and false negatives (FN) selected genes.

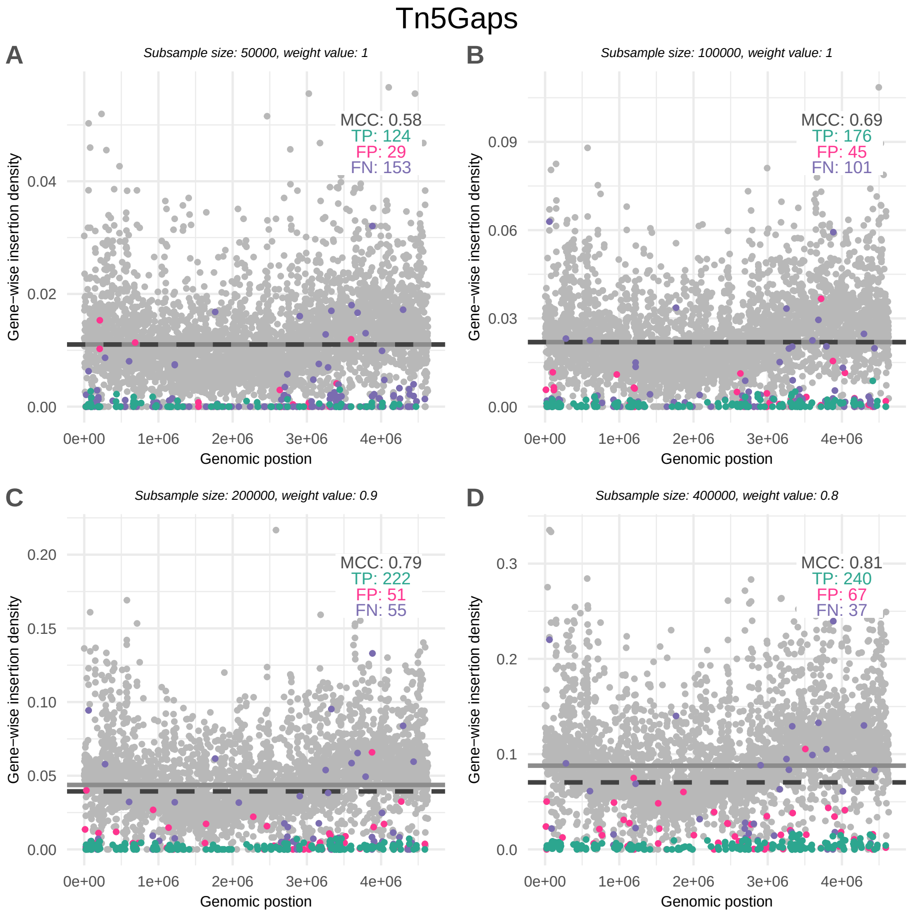


Figure S12: **Selected genes and gene-wise insertion densities for the Tn5Gaps approach**. The solid horizontal line shows genome-wide insertion density and the dashed horizontal line the weighted genome-wide insertion density giving the highest MCC. Green, pink and purple dots show the true positives (TP), false positives (FP) and false negatives (FN) selected genes.

### References

Tomoya Baba, Takeshi Ara, Miki Hasegawa, Yuki Takai, Yoshiko Okumura, Miki Baba, Kirill A Datsenko, Masaru Tomita, Barry L Wanner, and Hirotada Mori. Construction of escherichia coli k-12 in-frame, single-gene knockout mutants: the keio collection. *Molecular Systems Biology*, 2 (MSB4100050), January 2006.

Alissa Bleem, Ryo Kato, Zoe A. Kellermyer, Rui Katahira, Masahiro Miyamoto, Koh Niinuma, Naofumi Kamimura, Eiji Masai, and Gregg T. Beckham. Multiplexed fitness profiling by rb-tnseq elucidates pathways for lignin-related aromatic catabolism in sphingobium sp. syk-6. *Cell Reports*, 42(8):112847, August 2023.

Emily C. A. Goodall, Ashley Robinson, Iain G. Johnston, Sara Jabbari, Keith A. Turner, Adam F. Cunningham, Peter A. Lund, Jeffrey A. Cole, and Ian R. Henderson. The essential genome of escherichia coli k-12. *mBio*, 9(1), March 2018.

Yibing Ma, Mattia Pirolo, Bimal Jana, Viktor Hundtofte Mebus, and Luca Guardabassi. The intrinsic macrolide resistome of escherichia coli. *Antimicrobial Agents and Chemotherapy*, 68(8), August 2024.

Rabindra K. Mandal and Young M. Kwon. Global screening of salmonella enterica serovar typhimurium genes for desiccation survival. *Frontiers in Microbiology*, 8, sep 2017.

Kieran B. Pechter, Larry Gallagher, Harley Pyles, Colin S. Manoil, and Caroline S. Harwood. Essential genome of the metabolically versatile alphaproteobacterium rhodopseudomonas palustris. *Journal of Bacteriology*, 198(5):867–876, March 2016.

Kelly M. Wetmore, Morgan N. Price, Robert J. Waters, Jacob S. Lamson, Jennifer He, Cindi A. Hoover, Matthew J. Blow, James Bristow, Gareth Butland, Adam P. Arkin, and Adam Deutschbauer. Rapid quantification of mutant fitness in diverse bacteria by sequencing randomly bar-coded transposons. *mBio*, 6(3), July 2015.
