## Supplementary file 3 for "ConNIS and labeling instability: new statistical methods for improving the detection of essential genes in TraDIS libraries"

All R code is available under https://github.com/bips-hb/ConNIS_results and https://doi.org/10.5281/zenodo.16790977

**Setup for generating synthetic data**

For the generation of synthetic data the *Escherichia coli* K-12 strain MG1655 genome was used as a reference with respect for the length and position of genes. Based on the reported number of essential genes in real-world applications in the DEG 15 database (Luo et al., 2020), approximately 8% of the genes were randomly assigned as 'true essential genes'. Since essential genes can occur adjacent to each other across the genome, 100 clusters of essential genes were determined by drawing from a negative binomial distribution $NB(1, 0.3)$. These clusters of essential genes were then randomly distributed across the genome and each essential gene $j$ was characterized by a *minimum* length of an insertion-free sequence relative to its length $b_j$. The relative length of an insertion-free sequence was determined by sampling from a uniform distribution $\mathcal{U}(\lambda, 1)$, where $\lambda \in \{0.7, 0.75, 0.8, 0.85\}$, corresponding to *average* insertion-free regions of 85%, 87.5%, 90%, and 92.5% of the gene lengths, respectively.

A non-uniform distribution of IS along non-essential regions of the genome (cf. Zhang et al., 2021; Green et al., 2012; Mahmutovic et al., 2020; van Opijnen and Levin, 2020; Kimura et al., 2016; Manna et al., 2007), was attempted to be mimicked with the following two simulation schemes.

**Simulation scheme 1:** The regional IS densities across the genome should follow a sinusoidal shape. Therefore, an initial value $\frac{1}{b}$ was assigned to each base pair, where $b$ denotes the number of base pairs of the genome. Subsequently, all values were multiplied by a value from a wave function with a crest of 1.3 and a trough of 0.7 per cycle. The cycle length was set to 2 million base pairs. To introduce further variability, a random value from a normal distribution $\mathcal{N}(0, \frac{1}{3b})$ was added to each value. In essential genes, the values for base pairs to contain an IS were then set to zero if they lay within the essential part of the gene (i.e., the non-insertion sequence determined by $\mathcal{U}(\lambda, 1)$). For the remaining base pairs in non-essential parts of the genome, each value was normalized by dividing it by the sum of all values. This gave the final probability of each base pair containing an insertion site by chance.

**Simulation scheme 2:** All base pairs in non-essential regions of the genome were assigned an equal probability of containing an IS, except for genomic hot and cold spots. Within these spots, the probability of an observed IS was either elevated (hotspots) or reduced (coldspots). To generate the probabilities, each potential IS was initially assigned a value of $\frac{1}{b}$. Next, either 25 hotspots, 25 coldspots, 25 hot- and 25 coldspots, or no hot- and coldspots were randomly distributed across the genome. Hotspots and coldspots were defined as regions spanning 10,000 bp. The probability of an insertion at each base pair was scaled by factors of 5 (hot spots) and 0.1 (cold spots). Similar to simulation scheme 1, a random value from a normal distribution $\mathcal{N}(0, \frac{1}{3b})$ was added to each value to generate more variability. All values were normalized by dividing each value by the sum of all values to generate the final probabilities.

Using the probabilities of scheme 1 or 2 libraries of 50,000, 100,000, 200,000, and 400,000 IS were generated. Finally, either none or an extra 2% of IS were randomly added to the genome to address (technical) noise. These IS could appear anywhere in the genome, even in formerly insertion-free sequences of essential genes. Figure S1 (see Supplement 2) illustrates examples of our simulated data compared to real-world data, highlighting gene-wise insertion densities across the genome.

Hao Luo, Yan Lin, Tao Liu, Fei-Liao Lai, Chun-Ting Zhang, Feng Gao, and Ren Zhang. DEG 15, an update of the Database of Essential Genes that includes built-in analysis tools. *Nucleic Acids Research*, 49(D1):D677–D686, 10 2020.

Anel Mahmutovic, Pia Abel zur Wiesch, and Sören Abel. Selection or drift: The population biology underlying transposon insertion sequencing experiments. *Computational and Structural Biotechnology Journal*, 18:791–804, 2020.

Dipankar Manna, Steffen Porwollik, Michael McClelland, Ruimin Tan, and N. Patrick Higgins. Microarray analysis of mu transposition in salmonella enterica, serovar typhimurium: transposon exclusion by high-density dna binding proteins. *Molecular Microbiology*, 66(2):315–328, September 2007.

Tim van Opijnen and Henry L. Levin. Transposon insertion sequencing, a global measure of gene function. *Annual Review of Genetics*, 54(1):337–365, November 2020.

Houyu Zhang, Ting Lu, Shan Liu, Jianyu Yang, Guohuan Sun, Tao Cheng, Jin Xu, Fangyao Chen, and Kuangyu Yen. Comprehensive understanding of tn5 insertion preference improves transcription regulatory element identification. *NAR Genomics and Bioinformatics*, 3(4), October 2021.
