## Supplementary file 7 for "ConNIS and labeling instability: new statistical methods for improving the detection of essential genes in TraDIS libraries"

### Proof of concept for determining quasi-essential and fintness related genes using ConNIS and gene-labeling instability

This document presents a proof of concept for identifying quasi-essential genes and fitness effects using *ConNIS*. We do not, however, propose a full mathematical framework that incorporates a stochastic model for insertion sites or read counts over time and condition. Instead, the approach presented here is similar to previous work in that it heuristically compares gene essentiality or read counts between two fixed time points or conditions.

As an example, we use the work of *Schreier et al. (2023)* (https://doi.org/10.1073/pnas.2217200120). The authors measured the fitness of *Ruegeria pomeroyi* DSS-3 in a multispecies microbial environment. Their library consists of about 52,000 transposon mutants at baseline (t0, initial) and about 43,000 mutants after an 8-day growth period in monoculture (Rp, no other species). Since the genome length is about 4.1 million bp long, this library can be considered as a rather low-density transposon library, thus, making it challenging to avoid an inflation of false positives. We further compare a relative fitness score based on *ConNIS* with the relative fitness of the original work when other species are present (adjusted for the initial insertion sites).

```
# install.packages("devtools")
devtools::install_github("bips-hb/ConNIS")
```

First, we load the insertion positions in the initial library and in the Rp condition.

```
library(tidyverse)
library(gmp)
library(ConNIS)
library(readxl)
library(latex2exp)

source("functions.R")

# Organism: Ruegeria pomeroyi DSS-3
# Data & publication: https://www.pnas.org/doi/10.1073/pnas.2217200120

IS_raw <- readxl::read_xlsx("./DSS3/pnas.2217200120.sd02.xlsx", skip = 2)

IS_pos_initial <- IS_raw %>%
  mutate(readCount = InitialLibrary_1 +
                     InitialLibrary_2 +
                     InitialLibrary_3 +
                     InitialLibrary_4) %>%
  filter(readCount > 0) %>%
  select(genomePosition,readCount)

IS_pos_RP <- IS_raw %>%
  mutate(readCount = Rp_1 +
                     Rp_2 +
                     Rp_3 +
```

```
                    Rp_4) %>%
  # restrict to positions that were already present in the initial library
  filter(readCount > 0 &
           InitialLibrary_1 +
           InitialLibrary_2 +
           InitialLibrary_3 +
           InitialLibrary_4 > 0) %>%
  select(genomePosition,readCount)

gene_list <- read_tsv("./DSS3/R_pomeroyi_genes.tsv")
```

To account for non-uniform insertion densities, we use the instability function to determine a weighting value $w$ for *ConNIS* based on the initial library (un-comment code to run code).

```
### uncomment to run stability approach to determine a weighting factor for ConNIS

# connis_instabilities_initial <- ConNIS::instabilities(
#   ins.positions = IS_pos_initial$genomePosition,
#   gene.names    = gene_list$locus_tag,
#   method        = "ConNIS",
#   gene.starts   = gene_list$start,
#   gene.stops    = gene_list$end,
#   genome.length = 4109437,
#   d             = 0.5,   # fraction of original IS used in subsamples
#   m             = 500,   # number of subsamples
#   weights       = seq(0.05, 1, 0.05),
#   use.parallelization   = TRUE,
#   parallelization.type  = "mclapply",
#   numCores              = 25,
#   seed                  = 2025,
#   set.rng               = "L'Ecuyer-CMRG". # random number generator
# )
#
# saveRDS(
#   connis_instabilities_initial,
#   file = "results/connis_instabilities_initial_RuegeriaPomeroyiDSS3.RDS"
# )
```

From the sequence of instability values (see previous code block), we select the weight with the lowest instability.

```
connis_instabilities_initial <- readRDS(
  "results/connis_instabilities_initial_RuegeriaPomeroyiDSS3.RDS"
)

selected_w_initial <-
  min_instability(connis_instabilities_initial$weight_value,
                  connis_instabilities_initial$instability)
```

Next, we apply *ConNIS* with the selected weight to determine essential genes in the initial library (using Bonferroni correction for $\alpha$).

```
results_initial <- ConNIS::ConNIS(
  ins.positions = IS_pos_initial$genomePosition,
  gene.names    = gene_list$locus_tag,
  gene.starts   = gene_list$start,
```

```r
  gene.stops    = gene_list$end,
  genome.length = 4109437,
  weight        = selected_w_initial
)

alpha_star <- 0.05 / nrow(gene_list)  # Bonferroni-adjusted significance level
essential_initial <- gene_list$locus_tag[results_initial$p_value <= alpha_star]
```

We then run *ConNIS* for the Rp setting after the 8-day growth period, with a weight $w$ derived from the instability approach for this library.

```r
# load instability values for RP (just run the instability code above with IS from RP)
connis_instabilities_RP <- readRDS(
  "results/connis_instabilities_RP_RuegeriaPomeroyiDSS3.RDS"
)

selected_w_RP <-
  min_instability(connis_instabilities_RP$weight_value,
                  connis_instabilities_RP$instability)


results_RP <- ConNIS::ConNIS(
  ins.positions = IS_pos_RP$genomePosition,
  gene.names    = gene_list$locus_tag,
  gene.starts   = gene_list$start,
  gene.stops    = gene_list$end,
  genome.length = 4109437,
  weight        = selected_w_RP
)

essential_RP <- gene_list$locus_tag[results_RP$p_value <= alpha_star]
```

#### Quasi-essenitality

While previous work has often used a binary system to define "quasi-essential" genes after multiple generations (e.g. non-essential at $t_0$ and essential at $t_x$ with $t_x > t_0$; see *Hutchison et al. (2016)*, DOI: 10.1126/science.aad6253), we use the *Shannon information* to quantify how "surprised" we are to observe the biggest gap within a gene at a time point or under a condition. For a gene $g$ at time/condition $t$ we define

$$S_{g,t} := -\log_2(p_{g,t}),$$

where $p_{g,t}$ is the probability for the biggest gap determined by ConNIS. Applying a threshold

$$\tau = -\log_2(\alpha^*),$$

where $\alpha^* = \alpha_{\text{star}}$ is the Bonferroni-adjusted significance level to control the FWER, we classify genes as

- **essential** if $S_{g,t_0} \geq \tau$ and $S_{g,t_x} \geq \tau$,
- **quasi-essential** if $S_{g,t_0} < \tau$ and $S_{g,t_x} \geq \tau$,
- **non-essential** if $S_{g,t_0} < \tau$ and $S_{g,t_x} < \tau$.

For the rare cases where $S_{g,t_0} \geq \tau$ but $S_{g,t_x} < \tau$, we assume possible artifacts (e.g. due to noisy data or borderline p-values).

Setting $t_0 := t_{\text{initial}}$ and $t_x := t_{\text{Rp}}$, we obtain a classification and a scatter plot of $S_{\text{initial}}$ vs. $S_{\text{Rp}}$. Figure 1 shows the gene-wise *ConNIS* essentiality scores for *R. pomeroyi* in the initial library and after an 8-day growth period in monoculture (Rp). Quasi-essential genes occupy an intermediate region between consistently non-essential and consistently essential genes, illustrating that ConNIS supports a graded notion of essentiality rather than a strictly binary one. The black diagonal dashed line represents the separation of scores between the two methods where no difference in the *Shannon information* between the initial an the later time point is given. As expected, most points lie on the line or left of it (the latter indicating that the gaps are getting wider along time). A few points are right of the diagonal line, which could indicate that these gene are either beneficial. Points in the lower right quadrant are possible artifacts due to noise.

```r
tau <- -log2(alpha_star)

essential_data <- tibble(
  Gene      = gene_list$locus_tag,
  p_initial = results_initial$p_value,
  p_RP      = results_RP$p_value
)

essential_data <- essential_data %>%
  mutate(
    S_initial = -log2(p_initial),
    S_RP      = -log2(p_RP),
    essentiality = case_when(
      S_initial >= tau & S_RP >= tau ~ "essential",
      S_initial <  tau & S_RP >= tau ~ "quasi-essential",
      S_initial <  tau & S_RP <  tau ~ "non-essential",
      S_initial >= tau & S_RP <  tau ~ "possible artifact"
    )
  )

essential_data$essentiality[is.na(essential_data$essentiality)] <- "possible artifact"

ggplot(essential_data,
       aes(x = log(S_initial), y = log(S_RP), color = essentiality)) +
  geom_point(alpha = 0.6, size = 1.5) +
  geom_hline(yintercept = log(tau), color = "lightgrey") +
  geom_vline(xintercept = log(tau), color = "lightgrey") +
  geom_abline(slope = 1, color = "black", linetype="dashed") +
  scale_color_manual(values = c(
    "non-essential"     = "lightgreen",
    "quasi-essential"   = "orange",
    "essential"         = "violet",
    "possible artifact" = "lightblue"
  )) +
  theme_bw() +
  xlab(TeX("$\\log(S_{initial})$")) +
  ylab(TeX("$\\log(S_{Rp})"))+
  theme_dark()
```

Overall, our genes are classified as:

```r
table(essential_data$essentiality)
```

```
##
##         essential     non-essential possible artifact   quasi-essential
##               167              3896                 3               276
```

Since our instability approach and ConNIS favor rather sparse "models" in low density libraries (to reduce the otherwise inflation of false positives) the number of essential and quasi-essential labeled genes seems

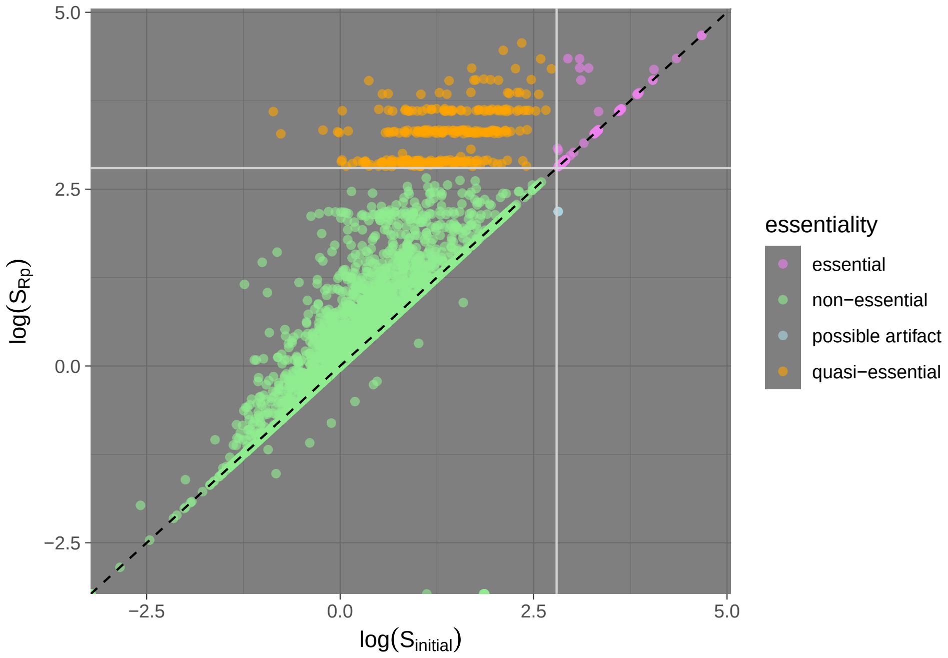


Figure 1: Essential, quasi-essential, non-essential and possible artifact genes based on ConNIS scores in the initial library and the Rp condition. The grey horizontal and vertical lines indicate the threshold corresponding to the Bonferroni-adjusted significance level. Genes are classified as non-essential (green), essential (violet), quasi-essential (orange; non-essential in the initial library but essential in Rp), or possible artifacts (blue; essential in the initial library but not in Rp).

reasonable.

#### Fitness scores

Next, we define a ConNIS-based fitness score and compare it with the fitness scores of the original work via correlation. Similar to the original work of *Schreier et al. (2023)* we do not compare the scores (in our case the *Shannon information*) directly between a community condition and RP to evaluate the relative gene fitness. Instead, the score of each gene with or without a community condition is first divided by the score of the initial library. Then, the relative gene fitness is calculated as the the $log_2$ of the ratio between RP and a community condition. For comparison, we load the fitness data of *Schreier et al. (2023)* and combine all to ione dataset.

```
# get insertion sites for the community conditions (restricted to positions
# present in the initial library)
IS_pos_RPplusV <- IS_raw %>%
  mutate(readCount = `Rp+V_1` +
                     `Rp+V_2` +
                     `Rp+V_3` +
                     `Rp+V_4`) %>%
  filter(readCount > 0 &
           InitialLibrary_1 +
           InitialLibrary_2 +
           InitialLibrary_3 +
           InitialLibrary_4 > 0) %>%
  select(genomePosition)
```

```r
IS_pos_RPplusM <- IS_raw %>%
  mutate(readCount = `Rp+M_1` +
                     `Rp+M_2` +
                     `Rp+M_3` +
                     `Rp+M_4`) %>%
  filter(readCount > 0 &
           InitialLibrary_1 +
           InitialLibrary_2 +
           InitialLibrary_3 +
           InitialLibrary_4 > 0) %>%
  select(genomePosition)

IS_pos_RPplusVandM <- IS_raw %>%
  mutate(readCount = `Rp+V+M_1` +
                     `Rp+V+M_2` +
                     `Rp+V+M_3` +
                     `Rp+V+M_4`) %>%
  filter(readCount > 0 &
           InitialLibrary_1 +
           InitialLibrary_2 +
           InitialLibrary_3 +
           InitialLibrary_4 > 0) %>%
  select(genomePosition)


# (we assume that the instability approach for the RP+V, RP+M and RP+V+M community conditions
# has been applied before; modify the code from above accordingly)

# load instability values for RP+V
connis_instabilities_RPplusV <- readRDS(
  "results/connis_instabilities_RPplusV_RuegeriaPomeroyiDSS3.RDS"
)

selected_w_RPplusV <-
  min_instability(connis_instabilities_RPplusV$weight_value,
                  connis_instabilities_RPplusV$instability)


## run ConNIS for the community conditions (using the same weight as for the initial library)
results_RPplusV <- ConNIS::ConNIS(
  ins.positions = IS_pos_RPplusV$genomePosition,
  gene.names    = gene_list$locus_tag,
  gene.starts   = gene_list$start,
  gene.stops    = gene_list$end,
  genome.length = 4109437,
  weight        = selected_w_RPplusV
)

# load instability values for RP+M
connis_instabilities_RPplusM <- readRDS(
  "results/connis_instabilities_RPplusM_RuegeriaPomeroyiDSS3.RDS"
)
```

```r
selected_w_RPplusM <-
  min_instability(connis_instabilities_RPplusM$weight_value,
                  connis_instabilities_RPplusM$instability)


results_RPplusM <- ConNIS::ConNIS(
  ins.positions = IS_pos_RPplusM$genomePosition,
  gene.names    = gene_list$locus_tag,
  gene.starts   = gene_list$start,
  gene.stops    = gene_list$end,
  genome.length = 4109437,
  weight        = selected_w_RPplusM
)

# load instability values for RP+V+M
connis_instabilities_RPplusVandM <- readRDS(
  "results/connis_instabilities_RPplusVandM_RuegeriaPomeroyiDSS3.RDS"
)

selected_w_RPplusVandM <-
  min_instability(connis_instabilities_RPplusVandM$weight_value,
                  connis_instabilities_RPplusVandM$instability)


results_RPplusVandM <- ConNIS::ConNIS(
  ins.positions = IS_pos_RPplusVandM$genomePosition,
  gene.names    = gene_list$locus_tag,
  gene.starts   = gene_list$start,
  gene.stops    = gene_list$end,
  genome.length = 4109437,
  weight        = selected_w_RPplusVandM
)

# load fitness data of Schreier et al. (2023)
fitness_data_original <-
  readxl::read_xlsx("./DSS3/pnas.2217200120.sd03.xlsx", skip = 3)

# convert fitness columns to numeric
for (i in 5:21) {
  fitness_data_original[, i] <- as.numeric(unlist(fitness_data_original[, i]))
}
rm(i)

# ConNIS-based scores and fitness proxies
# comparable to the original work  we use not the raw scores of each
# condition but rather the scores realtive to the initial scores.
fitness_data_connis <- tibble(
  Gene         = gene_list$locus_tag,
  p_initial    = results_initial$p_value,
  p_RP         = results_RP$p_value,
  S_initial    = -log2(p_initial + 1e-100),
  S_RP         = -log2(p_RP + 1e-100),
  w_RP         = S_RP - S_initial,
```

```r
  p_RPplusV    = results_RPplusV$p_value,
  S_RPplusV    = -log2(p_RPplusV + 1e-100),
  w_RPplusV    = S_RPplusV - S_initial,
  p_RPplusM    = results_RPplusM$p_value,
  S_RPplusM    = -log2(p_RPplusM + 1e-100),
  w_RPplusM    = S_RPplusM - S_initial,
  p_RPplusVandM = results_RPplusVandM$p_value,
  S_RPplusVandM = -log2(p_RPplusVandM + 1e-100),
  w_RPplusVandM = S_RPplusVandM - S_initial
)

# merge data for common genes
genes_for_comparison <- intersect(fitness_data_connis$Gene,
                                  fitness_data_original$Gene)

fitness_data_connis_reduced <- fitness_data_connis %>%
  filter(Gene %in% genes_for_comparison)

fitness_data_original_reduced <- fitness_data_original %>%
  filter(Gene %in% genes_for_comparison)

data_comparison <- right_join(
  x = fitness_data_connis_reduced,
  y = fitness_data_original_reduced,
  by = "Gene"
)

# remove any remaining rows with NA in relevant columns
data_comparison <- data_comparison[apply(data_comparison, 1, function(row) {
  !any(is.na(row))
}), ]
```

We now compare *ConNIS*-based relative fitness scores with the original fitness scores (community vs Rp) and define relative fitness difference as

$$\Delta(w_1, w_2) := w_{community} - w_{RP}$$

```r
# difference in w between communities and Rp; flip sign to be in correspondence with Schreier et al. (2
delta_w_RPplusV      <- -(data_comparison$w_RPplusV      - data_comparison$w_RP)
delta_w_RPplusM      <- -(data_comparison$w_RPplusM      - data_comparison$w_RP)
delta_w_RPplusVandM  <- -(data_comparison$w_RPplusVandM  - data_comparison$w_RP)

cor(delta_w_RPplusV,
    data_comparison$`log2_foldchange_Rp+V`,
    use = "complete.obs")
```

```
## [1] 0.5087618
```

```r
cor(delta_w_RPplusM,
    data_comparison$`log2_foldchange_Rp+M`,
    use = "complete.obs")
```

```
## [1] 0.4296997
```

```r
cor(delta_w_RPplusVandM,
    data_comparison$`log2_foldchange_Rp+V+M`,
    use = "complete.obs")
```

```
## [1] 0.5716215
```
